# saks-nf: A json solution for Nextflow pipeline construction

**DOI:** 10.1101/2022.09.19.508305

**Authors:** Xinming Zhuo, Nicholas Renzette, Gregory Omerza

## Abstract

To address the increasing complexity of data in scientific research, researchers have developed many workflow manager tools. Nextflow is one of the most widely used tools, enabling scalability and reproducibility of scientific workflows across various computational platforms through the implementation of domain-specific language (DSL) with a dataflow paradigm. We developed saks-nf, a JavaScript Object Notation (JSON) solution for constructing Nextflow pipelines. Our solution flattens the learning curve for Nextflow. Users can build and maintain a pipeline without prior programming experience by editing a JSON specification on any text editor. The saks-nf solution can be used to construct a multi-step pipeline with parallel and scatter-gather capacity in a computing platform of choice, such as a local machine, a high-performance cluster, or cloud computing. This solution allows users to focus on analysis, thereby increasing productivity.

## 1. Introduction

With the rapid and broad adaptation of high-throughput technology, the volume and complexity of data increases exponentially, resulting in the need of complex analysis pipelines with an extensive array of software. While these analyses enable rapid scientific advances, they typically consist of multi-step processes and are hindered by variability in computational environments and ambiguities in parameters or software versions, thus limiting scalability and reproducibility. Many institutes and research groups are developing various workflow managers to address these issues. In data-centric industries, many companies have successfully deployed their home-brew workflow managers, such as the cadence by Uber (https://github.com/uber/cadence) and luigi by Spotify (https://github.com/spotify/luigi) [1]. In academia, especially in biomedical research, multiple workflow managers also have been adapted, such as the Galaxy Project [2], Cromwell+WDL [3], Snakemake [4, 5] and Nextflow [6, 7]. These tools assist bioinformaticians in many stages of the analysis pipeline lifecycle, including development, testing, and maintenance in order to enhance the portability and reproducibility.

Nextflow is one of the most widely used workflow managers [1]. It uses a domain-specific language (DSL) with a dataflow paradigm that enables rapid pipeline development compatible with any scripting language [6]. It is highly portable with multi-platform (OS and conda) support and container management (Docker, Singularity, etc.) [8, 9]. It can achieve scalability beyond the local infrastructure by providing built-in support for high-performance computing environments (such as SLURM, Univa Grid Engine, PBS/Torque, LSF ad HTCondor) and cloud computing services (such as AWS, Azure Cloud, and Google Cloud). Nextflow also allows reentry when the pipeline is disrupted, enabling users to run a pipeline from its last successfully executed step rather than from the beginning to save significant time and computing resources. Nextflow integrates with software repositories (GitHub) to enable robust version control and seamless pipeline sharing. Finally, Nextflow has a large active community and hosts a community curated pipeline repository, nf-core [10], which provides many ready-to-use and well-maintained pipelines.

Nextflow is well-suited for bioinformaticians with prior programming experience. It is written with an extension of the Groovy programing language and lacks a user-friendly workflow specification (Cromwell + WDL) or graphical user interface (Galaxy), which makes the initial learning curve steep [1]. To flatten the learning curve of Nextflow, a human readable/writable specification with a lightweight data-interchange format is desirable, such as JavaScript Object Notation (JSON) [11] or Yet Another Markup Language (YAML) (https://yaml.org). Here we present “Swiss Army Knife Solution for nextflow” (saks-nf), a JSON Application Programming Interface (API) for workflow specification in Nextflow that helps users compose a basic Nextflow pipeline without prior workflow experience. saks-nf also fully integrates with GitHub practice, enabling easy pipeline sharing and good reproducibility. It employs Nextflow’s strong support for multi-platform, container, and cloud computing and implements process parallelization and scatter-gather with minimal effort. This JSON-API follows the guidelines of the shared convention (https://jsonapi.org/), assists users in increasing productivity, and takes advantage of generalized tooling, so the users can focus on what matters most: analysis.

## 2. Materials and Methods

saks-nf is written in shell, json and groovy scripts and follows the instructions and guidelines of Nextflow. The saks-nf was tested in Ubuntu 16.04 and later versions of Linux with jq (v1.6 or later), sed (GNU sed v 4.7 or later), awk (GNU Awk 5.0.1 or later). The Nextflow pipeline generated by sak-nf is compatible with Nextflow (v21.04 or later). The workflow is also compatible with git for maintenance and sharing (git version 2.25 or later).

## 3. Results

### 3.1 Constructing a Nextflow pipeline

saks-nf is integrated with GitHub as a repository (https://github.com/xmzhuo/saks-nf). The main component is a shell script (run.sh), three Nextflow script templates (template.nf, modules/sak.nf, and modules/sak_docker.nf), and multiple config files (nextflow.config and multiple platform-specific config files in the config directory). Two additional directories (bin and data) contain placeholder files. The sak_data directory provides various example input files, such as bed files, dict files, and multiple json files and is included for training/practicing purposes. We also provide directory sak_example_output to host example output files.

To run saks-nf, the user must confirm jq and nextflow are appropriately installed, then run the shell script run.sh (bash saks-nf/run.sh -i example.json). With an “-o” argument, the user can designate a specific location and name for the composed Nextflow pipeline; otherwise, the new workflow will be named after the json file at the exact location. Users also can force overwrite an existing folder with the desired name with an “-f” argument. If users prefer to run the new workflow immediately, they can use the “-r” argument. It is recommended to use “nextflow run” to execute a new workflow only after manually inspecting the composed workflow, especially for HPC or cloud computing environments.

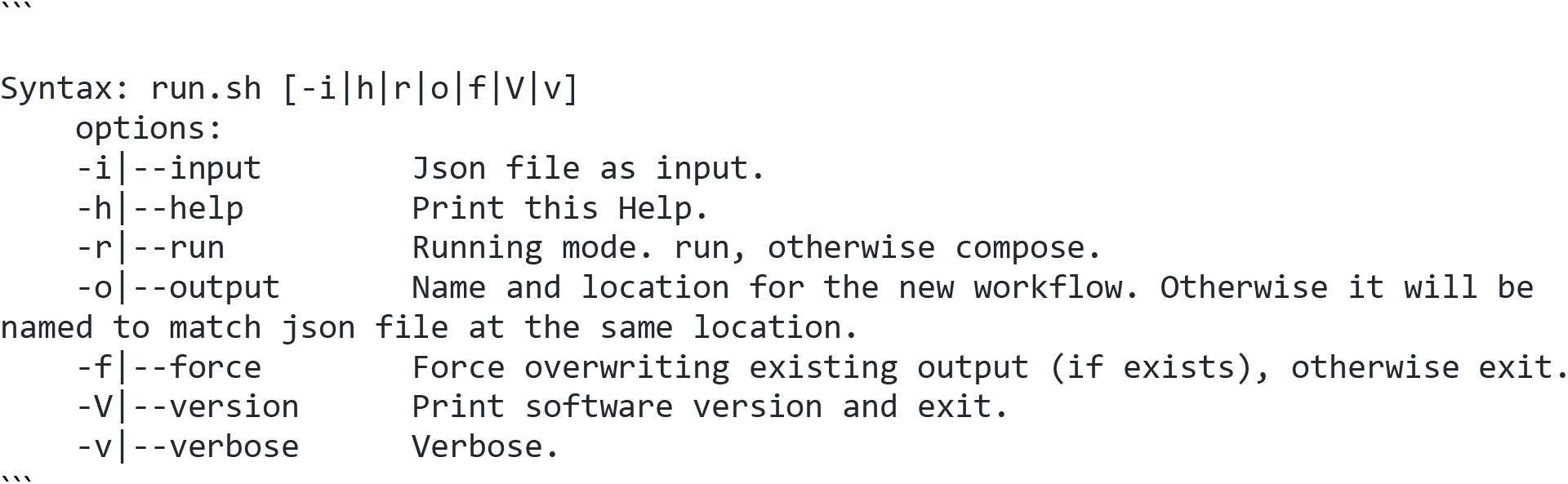

Once the new workflow is constructed with example.json, it will be named after the json file, example-nf. The workflow consists of main.nf, nextflow.config, modules, and config folders. The modules folders contain two nextflow scripts for each process (one with container, and other one without container). The config folders contain config files for the platform of choice, which may require users to provide their credentials for specific services by editing the specific config files. Additional bin and data folders are merely placeholders unless users prefer to make further advanced customization. After inspecting/editing the workflow, users can kick off the analysis with “nextflow run /path/to/example-nf”.

### 3.2 Structure of a JSON input

Users need to compose a Nextflow workflow with a json file, which is a collection of name/value pairs [11]. The minimum JSON input for saks-nf contains five elements, “title”, “profile”, “workdir”, “reportdir” and “process” (Figure 1). Users can add additional elements they need, such as the “description”. The “title” pairs with a string for the workflow’s name. The “profile” is the string value for the config file names in the config folder, such as “standard” for the native local environment and “Azure” for Microsoft Azure cloud computing. The “workdir” designates a local or cloud location for storing temporary Nextflow files, which can be used for workflow reentrancy or troubleshooting. The “reportdir” is the designated location for an html workflow execution report. The “process” is the element for constructing a workflow of single or multiple steps in the form of objects.

**Figure 1.**
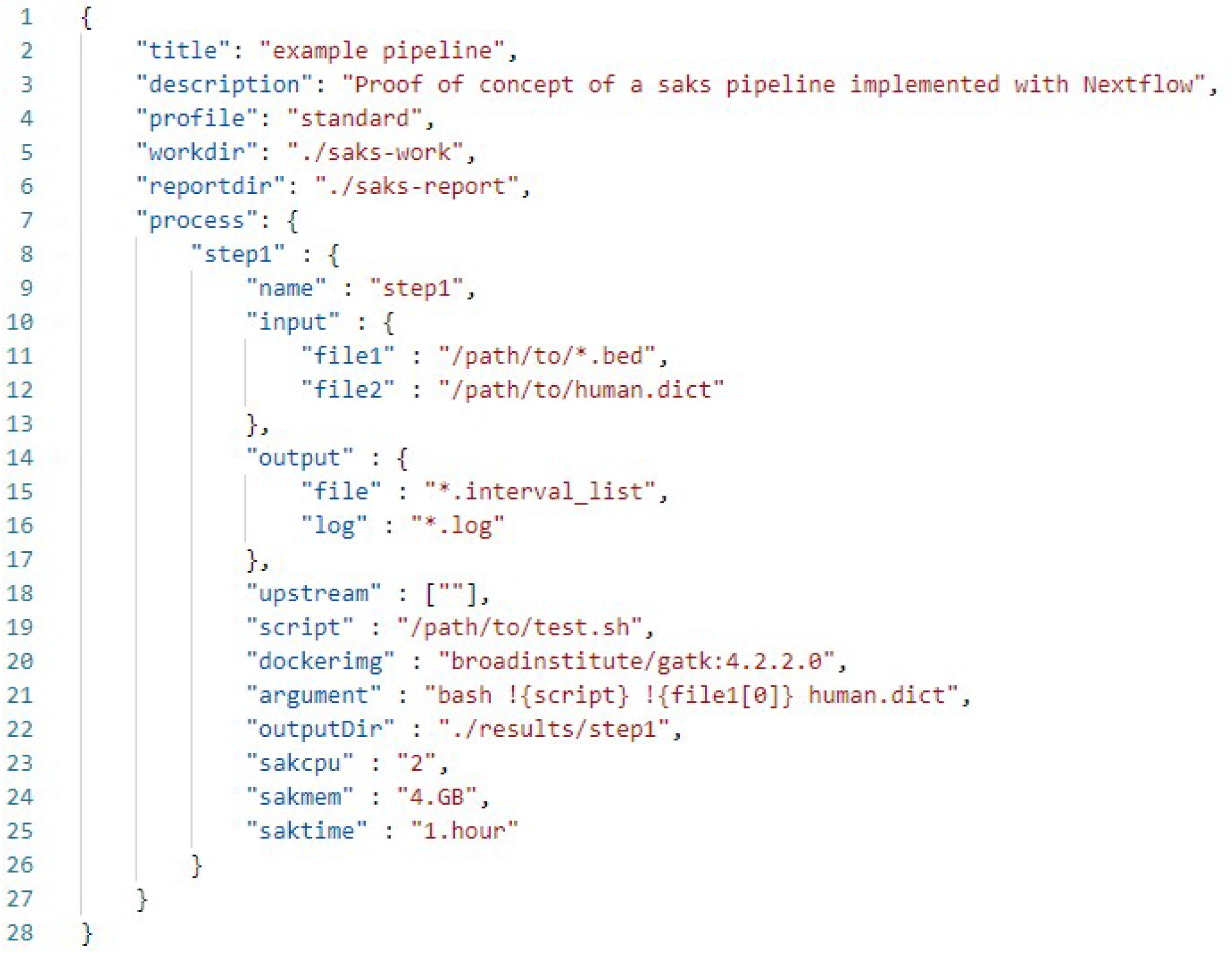
Example of a saks-nf json file. A minimum saks-nf json contains a collection of name/value pairs to specify a Nextflow workflow. The names are colored in blue and the values in red.

Each member of the “process” represents a specific step in a workflow. Therefore, every step has multiple elements to instruct Nextflow: the name, the computing environment, input, output, and command for the step. The step can have a customized name with a string value that matches the element “name”. The user then defines the “input” and “output” with objects. Each object can be assigned a customized name and string value pair that is compatible with regular expressions. For example, the “file1”: “/path/to/*.bed” for fetching all “bed” files in the path of ““/path/to/”. If this process depends on an upstream “step0” output “file”, the user can define the “upstream” in the form of an array ([“step0.file”]). Users can use an “argument” for a short and straightforward one-line shell script for simple execution following Nextflow convention (echo a process-specific variable with “!”, a customized variable with “$”). In this example, we only use the first file in “file1”. For more complicated execution, the user can provide a path to a script (such as shell, R, Python) that is supported by the designated environment. If a docker image is needed, users can specify the “dockerimg”, otherwise the value can be blank (“”). The user can assign the “outputDir”, “sakcpu”, “sakmem”, “saktime”, for output path, CPU allowance, allocate memory and timeout policy respectively.

### 3.3 Example: Simple two-step pipeline

The sak_data/example_io.json (https://github.com/xmzhuo/saks-nf/blob/main/sak_data/example_io.json) demonstrates how to compose a simple two-process workflow (sak_example_output/standard/results) (Figure 2A). The first process, bed2interval, uses GATK from a public docker image, “broadinstitute/gatk:4.2.2.0”, to convert a bed file to an interval_list [12]. To demonstrate the usage of a “script” file and process-specific variable, we write “gatk BedToIntervalList” command into a shell script, “test.sh”. In the “argument”, the process-specific variable “!{script}” points to the “test.sh” script. We request two outputs from this process “file”: “*.interval_list” and “log”: “*.log”. The interval_list is the product of running the “script”. saks-nf also produces the running argument/script log and appends the md5sum value of the output files to the end of log.

**Figure 2.**
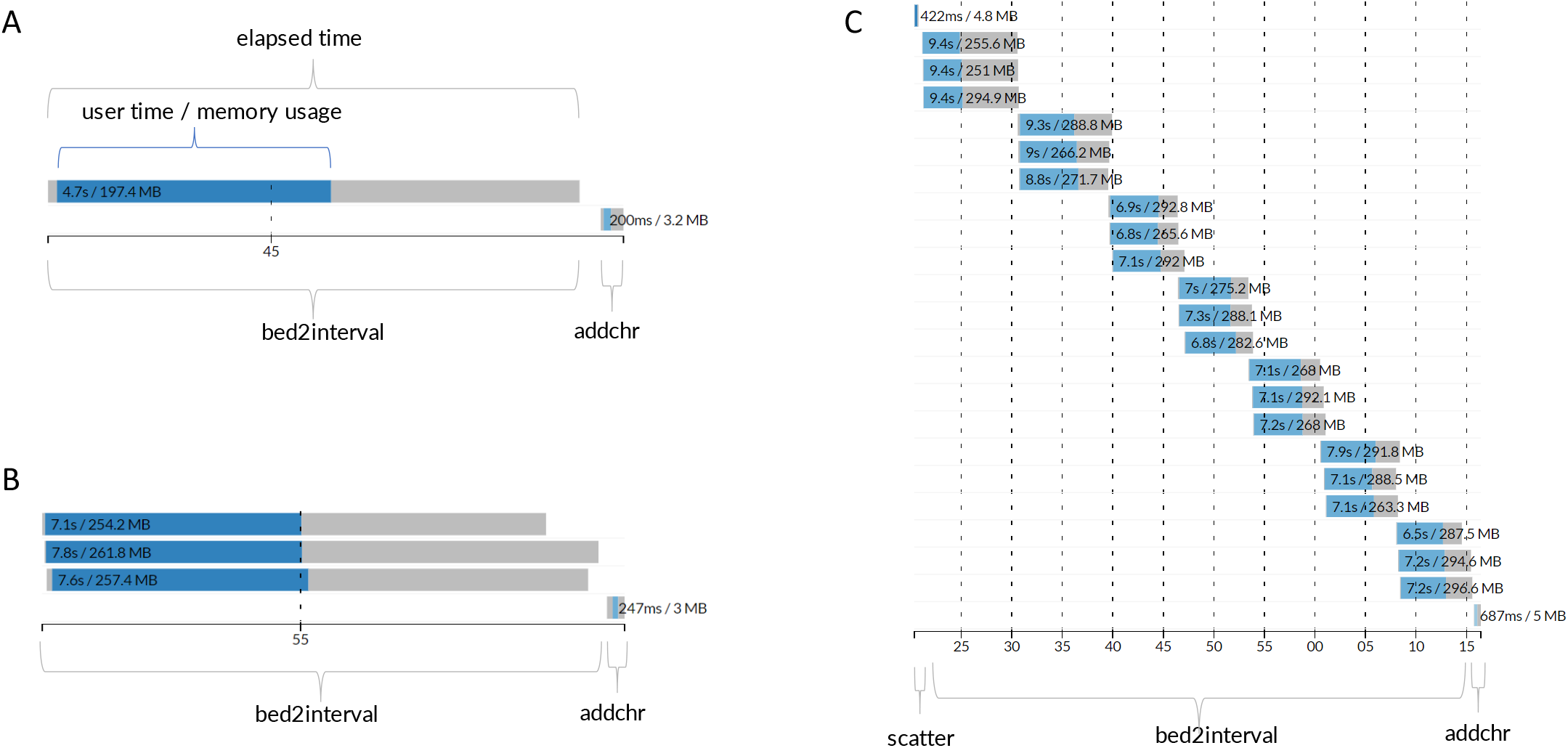
Overview of Using saks-nf to build Nextflow pipeline. (A) Example of using a json file to generate a Nextflow pipeline with two step analysis. The pipeline can be executed on different platforms: local, HPC and cloud. All the steps of the pipeline can be containerized to enhance reproducibility and portability. The output and report for the pipeline will be stored in a designated location. The same approach can be used for (B) a Nextflow pipeline with parallelized analysis, and (C) a Nextflow pipeline with scatter-gather analysis. In this example, only containerized steps are presented.

The second process, “addchr”, adds ‘chr’ string to the beginning of every line in the test.bed.interval_list from the last step. This step runs in a native bash shell environment, so a docker image is not needed. The “file” output from bed2interval were injected into the “upstream”: [“bed2interval.file”]. The “argument” demonstrates that the saks-nf can handle customized variable, $file, in a one-line bash script with piping. At the end, the final product “test.bed.chr.interval_list” was stored in the destination, “./results/addchr”. The timeline of one run with this workflow is shown in Figure 3A. The actual time of each step, including user time (blue) and elapsed time (blue and grey), are plotted accordingly.

**Figure 3.**
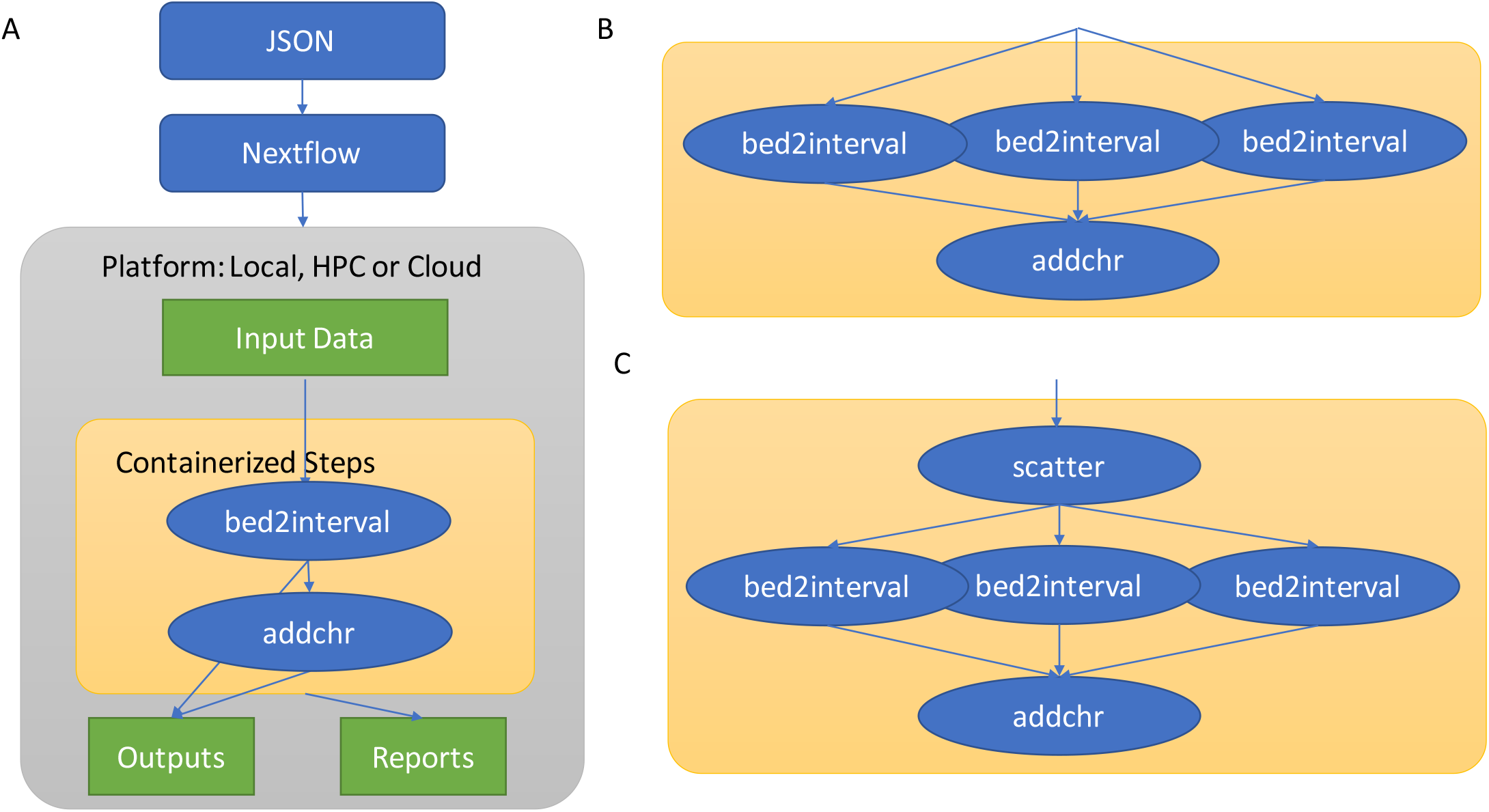
Timeline report of Nextflow pipeline. Timeline for (A) a standard two step pipeline, (B) a pipeline with a parallelized first step, and (C) a pipeline with scatter-gather structure. The x-axis represents total elapsed time (in seconds) for running each pipeline. Blue bars represent actual user time for each user step. Grey bars represent elapsed time for each user/step. All three timelines are pipelines running on a local machine with limited CPU cores which affects the maximum instances that can be scattered.

### 3.4 Example: Multi-platform and cloud-ready

saks-nf can compose Nextflow workflows useable on HPC or cloud environments with a specific “profile”. The users need to modify configuration files in config folders accordingly. We have provided some template config files for several different scenarios, such as slurm, AWS and Azure. For example, if the user would like to run the workflow on HPC with slurm [13], they can assign “slurm” to “profile” and confirm the options meet their HPC’s policy. Depending on the configuration of the HPC, users may need to compose the workflow first then use sbatch to run the workflow. If the user would like to run the workflow on Azure, the user needs to add their Azure credentials (storage account and batch account), assign “azure” to “profile”, and storage location to “outputDir”. They also need to check if the “vmType” meets each step’s needs and add a process-specific “queue” value accordingly. In the case of AWS, users need to add their credentials to the config file and go to the AWS website to create/check specific queues that meet the demand of each step and assign the “queue” value accordingly. We prefer a docker image with a known configuration for cloud computing to fulfill the dependencies in the process (as shown in the “addchr” process in the json for Azure) (https://github.com/xmzhuo/saks-nf/blob/main/sak_data/example_az.json).

### 3.5 Example: Parallelization and Scatter-Gather

The example_io-parallel.json (https://github.com/xmzhuo/saks-nf/blob/main/sak_data/example_io-parallel.json) is a modified version of the previous json file to demonstrate how to compose a Nextflow workflow with parallelization (Figure 2B). In this case, the input has six bed files (4.small.bed, 4.large.bed, 5.small.bed, 5.large.bed, 6.small.bed and 6.large.bed). The new workflow converts three pairs (6) of bed files to interval_llist in parallel and adds ‘chr’ to the beginning of each line of the interval_list files. In the first process, the six bed files were paired with ‘large’ and ‘small’ according to the instruction in “inputpairing”, “bed”: [“large”,”small”]. For example, “4.large.bed” and “4.small.bed” were paired and grouped into one channel. Thus, three channels were formed and fed into the bed2interval process parallelly. After the three bed2interval processes were finished, the output files were collected and triggered initiation by the “addchr” process. The example nextflow script and result can be found in “sak_example_output/parallel”. The timeline of each step is shown in Figure 3B.

The objects in “inputpairing” in a specific process instruct parallelization to Nextflow. If users prefer to parallelize the bed2interval with six channels rather than above mentioned three channels, they can use “bed”: [“bed”], thus each bed file can form an independent channel and six bed2interval processes will be launched parallelly. If parallelization is not necessary, the user can drop the “inputpairing” element, and the workflow will form a channel containing all bed files.

The parallelization not only applies to “input” but also to “upstream”, which allows scatter-gather. The example_io-scattergather.json (https://github.com/xmzhuo/saks-nf/blob/main/sak_data/example_io-scattergather.json) demonstrates a simplified example of scatter-gather with saks-nf (Figure 2C). It starts with a “scatter” process that splits a bed file by chromosome followed by a “bed2interval” process, which forms a channel for each chromosome bed by assigning “scatter_file”: [“bed”] in the “upstreampairing”. Then, this process uses a GATK tool inside a docker image, “broadinstitute/gatk:4.2.2.0”, to convert each chromosome bed file to interval_list in parallel. The third process collects all the interval_list from all “bed2interval” processes and adds ‘chr’ to the chromosome in interval_list. The example nextflow script and result can be found in “sak_example_output/scattergather”. The timeline of scatter-gather step is shown in Figure 3C. Since the number of parallel processes is limited by the available CPU and memory of the user’s computing environment. In this example the workflow run is this example only allocated three processes running in parallel.

## 4. Discussion and Conclusions

There is an increasing need for open and reproducible research practices from scientific societies [14]. Workflow managers have become essential for bioinformatic studies to increase the reproducibility and impact of scientific research [1]. As one of the widely accepted workflow managers, Nextflow has many valuable features, such as multi-platform support, native task, streaming processing, dynamic branch evaluation, code sharing integration, workflow versioning, automatic error failover, and DAG rendering. These features enhance reproducibility by simplifying robust and complex analysis implementation besides optimizing resource management. These advantages are steppingstones for achieving FAIR (findable, accessible, interoperable, and reusable) analysis and allowing collaboration with large-scale studies in science [15]. Another attractive feature of workflow managers is that many ready-to-use pipelines can be found in the public repository, such as nf-core, enabling many researchers to perform complex analyses without prior programming experience.

Bioinformatics software and analyses are evolving at lightning speed, and many new software packages and analysis solutions may not be available in the form of a ready-to-use pipeline. Our saks-nf provides a JSON solution to build Nextflow pipelines without prior knowledge of Groovy and DSL, which significantly flattens the steep learning curve for implementation. With the JSON format specification, saks-nf allows the separation of specification and workflow generation. The user can create and edit the specification file with any text editor and introduce additional elements for versioning or descriptions, enhancing the readability and portability of the workflow. Furthermore, it simplifies the process of building complex multiple process workflows with parallelization and scatter-gather. saks-nf also takes advantage of multi-platform support in Nextflow; the user can easily equip with docker or local executer and deploy across various platforms, such as local pc, HPC (slurm), or cloud with supplied template config files. In addition, users can configure the process’s requirement, such as cpu, mem, and timeout policy, with a minor updates of the json file. Finally, saks-nf has a built-in function to generate a log file with md5sum value. The latest version of saks-nf is hosted on Github (xmzhuo/saks-nf) with examples and brief tutorials. We anticipate it will be a valuable resource to help users without prior experience in constructing their Nextflow workflows.

## Acknowledgments

This work is supported by internal clinical research project in CLIA Laboratory in The Jackson Laboratory for Genomic Medicines.

## References

1. Wratten, L., A. Wilm, and J. Göke, Reproducible, scalable, and shareable analysis pipelines with bioinformatics workflow managers. Nat Methods, 2021. 18(10): p. 1161–1168.

2. Goecks, J., et al., Galaxy: a comprehensive approach for supporting accessible, reproducible, and transparent computational research in the life sciences. Genome Biol, 2010. 11(8): p. R86.

3. Voss, K., G. Van der Auwera, and J. Gentry, Full-stack genomics pipelining with GATK4 + WDL + Cromwell. F1000Res, 2017. 6.

4. Koster, J. and S. Rahmann, Snakemake-a scalable bioinformatics workflow engine. Bioinformatics, 2018. 34(20): p. 3600.

5. Koster, J. and S. Rahmann, Snakemake--a scalable bioinformatics workflow engine. Bioinformatics, 2012. 28(19): p. 2520–2.

6. Di Tommaso, P., et al., Nextflow enables reproducible computational workflows. Nat Biotechnol, 2017. 35(4): p. 316–319.

7. Strozzi, F., et al., Scalable Workflows and Reproducible Data Analysis for Genomics. Methods Mol Biol, 2019. 1910: p. 723–745.

8. Kurtzer, G.M., V. Sochat, and M.W. Bauer, Singularity: Scientific containers for mobility of compute. PLoS One, 2017. 12(5): p. e0177459.

9. Merkel, D., Docker: lightweight linux containers for consistent development and deployment. Linux journal, 2014. 239: p. 2.

10. Ewels, P.A., et al., The nf-core framework for community-curated bioinformatics pipelines. Nat Biotechnol, 2020. 38(3): p. 276–278.

11. Pezoa, F., et al., Foundations of JSON schema, in Proceedings of the 25th International Conference on World Wide Web., I.W.W.W.C.S. Committee, Editor. 2016. p. 263–273.

12. McKenna, A., et al., The Genome Analysis Toolkit: a MapReduce framework for analyzing next-generation DNA sequencing data. Genome Res, 2010. 20(9): p. 1297–303.

13. Andy, B., A. YooMorris, and J. Grondona, SLURM: Simple Linux Utility for Resource Management, in Job Scheduling Strategies for Parallel Processing, D. Feitelson, L. Rudolph, and U. Schwiegelshohn, Editors. 2003, Springer. p. 17.

14. Auer, S., et al., Science Forum: A community-led initiative for training in reproducible research. eLife, 2021(10).

15. Wilkinson, M., Dumontier, M., Aalbersberg, I. et al., The FAIR guiding principles for scientific data management and stewardship. Scientific Data, 2016. 3: p. 160018.

